# Proteomics reveals profound metabolic changes in the alcohol use disorder brain

**DOI:** 10.1101/447912

**Authors:** Charmaine Enculescu, Edward D. Kerr, K. Y. Benjamin Yeo, Peter R. Dodd, Gerhard Schenk, Marina R. S. Fortes, Benjamin L. Schulz

## Abstract

Changes in brain metabolism are a hallmark of Alcohol Use Disorder (AUD). Determining how AUD changes the brain proteome is critical for understanding the effects of alcohol consumption on biochemical processes in the brain. We used data-independent acquisition mass spectrometry proteomics to study differences in the abundance of proteins associated with AUD in pre-frontal lobe and motor cortex from autopsy brain. AUD had a substantial effect on the overall brain proteome exceeding the inherent differences between brain regions. Proteins associated with glycolysis, trafficking, the cytoskeleton, and excitotoxicity were altered in abundance in AUD. We observed extensive changes in the abundance of key metabolic enzymes, consistent with a switch from glucose to acetate utilization in the AUD brain. We propose that metabolic adaptations allowing efficient acetate utilization contribute to ethanol dependence in AUD.

## Introduction

The effects of alcoholism are an ongoing risk to both mental and physical health, and to the wellbeing of society. The World Health Organisation estimates that 76.3 million people worldwide are diagnosable with alcohol use disorder (AUD) ^1^, and it is estimated that the dangerous consumption of alcohol results in 2.5 million deaths worldwide every year and is the third leading risk factor for poor health globally. A more detailed understanding of the causes and consequences of AUD is relevant to mitigating its impact on society.

The detrimental cognitive effects caused by chronic alcohol consumption have been well documented ^2,3^ However, the underlying molecular mechanisms and neurochemistry of alcoholism remain elusive. In this context, systems biology has been a powerful tool to investigate the molecular changes in the brain associated with AUD. Changes in brain gene expression have been studied with DNA microarrays ^4^ and RNAseq ^5,6^, which found a multitude of genes and pathways that were associated with tolerance, dependence, and neurotoxicity in chronic alcohol abuse. In particular, there was a marked and coordinated reduction in mRNAs coding for myelin proteins in the frontal lobe in AUD patients, suggesting that chronic alcohol abuse either directly or indirectly affects myelin-related genes, or that a loss of expression may be due to oligodendrocyte susceptibility to ethanol ^4^. This is consistent with alcohol-related neuropathology, where demyelination is observed in the pre-frontal lobe^7,8^.

Protein abundance does not necessarily correlate with transcript expression ^9^, especially at a tissue level in a complex and heterogeneous organ such as the brain, which has a multitude of distinct cell types ^10-12^. It is recognized that the brain and circulating proteome may therefore also help identify risk factors, biomarkers, and underlying causes of AUD ^13^. Previous proteomic studies of the brain of human alcoholics have used two-dimensional gel-electrophoresis and mass-spectrometry ^14-16^. One study of the dorsolateral pre-frontal lobe found a lower abundance of thiamine-dependent proteins, as well as enzymes involved in glycolysis and the TCA cycle in alcoholic patients^16^. A more recent study identified reduced abundance of creatinine kinase chain B, NADH_2_ ubiquinone, fructose-bisphosphate aldolase C, and glyceraldehyde-3-phosphate dehydrogenase, proteins central to cellular metabolism^14^. While these studies suggest that AUD is associated with diverse changes to the brain proteome, this has not been tested with modern mass spectrometry proteomics. Here, we used sequential window acquisition of all theoretical mass spectra (SWATH-MS) proteomics to mine the deep brain proteome and identify proteomic correlates of AUD.

## Materials and Methods

### Tissue collection and storage

All brain tissue samples were provided by the Queensland Brain Bank (QBB), School of Chemistry and Molecular Biosciences, The University of Queensland, with informed consent from the next of kin. QBB is the Brisbane node of the Australian Brain Bank Network, which is supported by the National Health and Medical Research Council. Post-mortem histopathological examination was conducted by qualified pathologists, and tissue samples were treated as previously described^17^. The Medical Research Ethics Committee of The University of Queensland approved the study (Clearance N° 0000105).

### Case selection

Clinic information including age, post-mortem interval, brain weight, and cause of death were retrieved from the QBB. Tissue from pathologically affected pre-frontal lobe and relatively spared motor cortex were sectioned in 0.5 g pieces. AUD was defined by National Health and Medical Research Council/WHO criteria as individuals who consumed greater than 80 g of ethanol per day for most of their adult lives. Controls either did not consume alcohol or were social drinkers who consumed less than 20 g of ethanol per day on average. Five males were selected for control cases and six males for the AUD cases, all from North-European descent. Age, post-mortem interval (PMI), and brain weight were matched as closely as possible in accordance with brain and case availabilities (Table 1). Alcoholic subjects were confirmed to not have Wernicke-Korsakoff syndrome^3^.

**Table 1.**
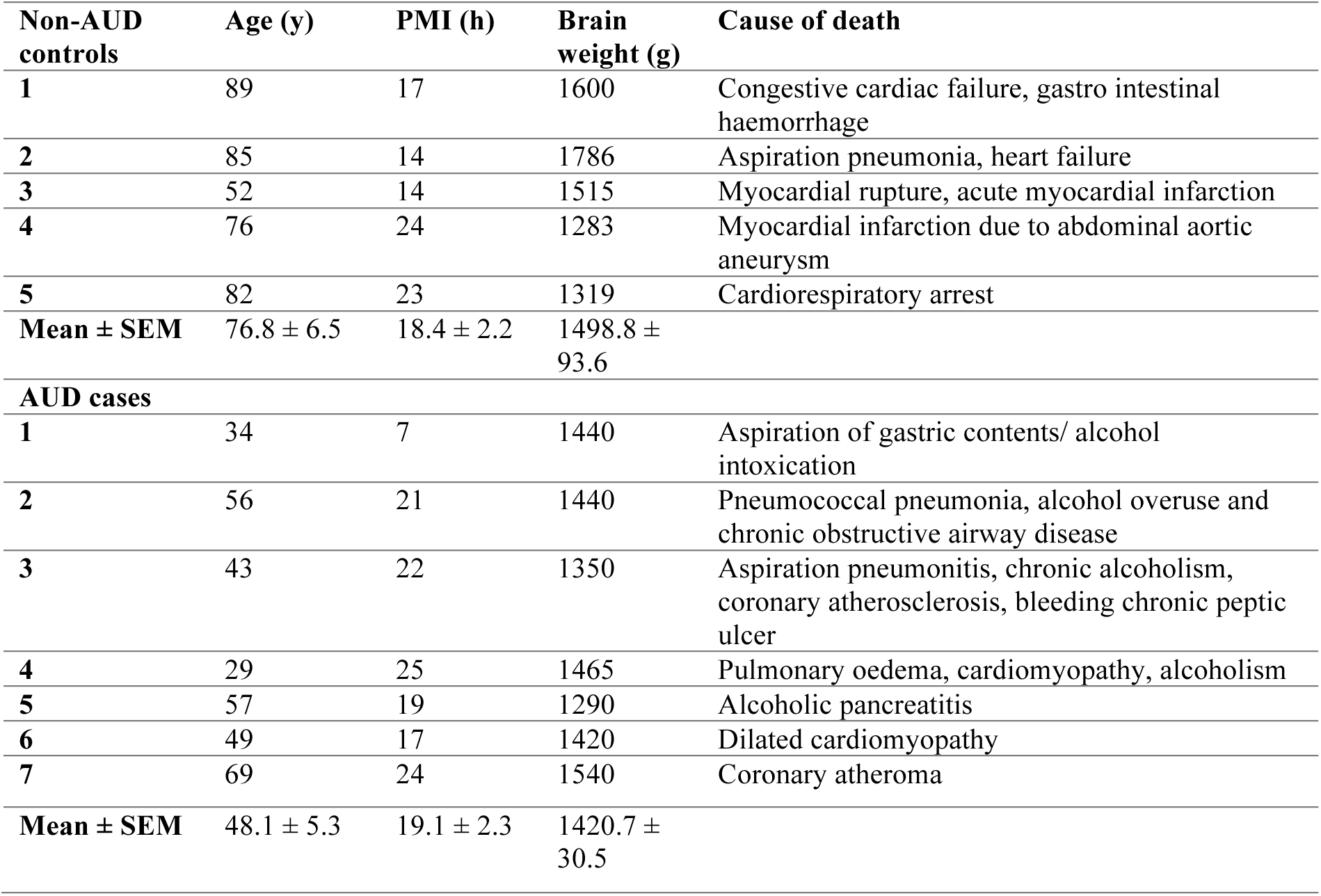
Case information.

### Whole tissue protein extraction

Whole tissue soluble protein extractions were prepared from the dorsolateral prefrontal gyrus of the pre-frontal lobe and the primary motor cortex of controls and alcoholic cases. The sectioned tissue pieces (0.5 g) were rapidly thawed in 0.32 M sucrose at 37 °C. Tissue was then immediately cooled to 4 °C on ice, weighed and homogenised in 10 volumes of ice-cold distilled water with a motor-driven Teflon-glass homogeniser. The homogenate was centrifuged at 1500 rcf for 5 min at 4 °C. Protein concentration was estimated via Lowry assay and adjusted to 3 mg/mL.

### Sample preparation

Samples were denatured and reduced/alkylated essentially as previously described ^18,19^. Briefly, samples containing approximately 100 μg of total protein were each suspended in 10 volumes of 50 mM TrisHCl buffer pH 8, 10 mM DTT, and 6 M guanidine HCl and incubated at 30 °C for 30 min with shaking. Cysteines were alkylated by addition of acrylamide to a final concentration of 30 mM and incubation at 30 °C for 1 h with shaking, and excess acrylamide was then quenched by addition of DTT to an additional final concentration of 10 mM. Samples were clarified by centrifugation at 18000 rcf for 10 min at room temperature. The supernatant was collected and processed with Filter Assisted Sample Preparation (FASP) ^20^ using a 10 kDa cut-off Amicon ultra 0.5 mL filter (EMD Millipore, Billerica, MA, USA) by centrifugation at 14000 rcf for 20 min at room temperature. 200 μL of 50 mM ammonium acetate was added to the filters, samples were further centrifuged at 14000 rcf for 20 min, and were then diluted with 130 μL of 50 mM ammonium acetate. Trypsin (proteomics grade, Sigma) was added in a 1:100 (enzyme:protein) ratio and the mixture was incubated overnight at 37 °C. The trypsin-digested peptides were then eluted by centrifugation at 14000 rcf for 20 min. Columns were washed with 50 μL of 0.5 M NaCl and centrifuged at 14000 rcf for 20 min. Trypsin-digested peptides were desalted with C18 ZipTips (Millipore) according to the manufacturer’s instructions. A pooled sample totalling approximately 40 μg peptides with a total volume of 500 μL in 0.1% formic acid was subjected to high pH reverse-phase fractionation. Peptides were applied to a 50 mg tC18 Sep-Pak (Waters), washed with 500 μL ddH_2_O, eluted in 8 separate 500 μL fractions of acetonitrile (5%, 7.5%, 10%, 12.5%, 15%, 17.5%, 20%, and 50%) in 0.1% triethylamine, lyophilised, and resuspended in 0.1% formic acid.

### Mass spectrometry proteomics

Peptides were identified by LC-MS/MS using a Prominence nanoLC system (Shimadzu) and a TripleTof 5600 mass spectrometer with a Nanospray III interface (SCIEX) essentially as previously described ^21^. Approximately 2 μg of peptides were desalted on an Agilent C18 trap (300 Å pore size, 5 μm particle size, 0.3 mm i.d. × 5 mm) at a flow rate of 30 μL/min for 3 min and then separated on a Vydac EVEREST reversed-phase C18 HPLC column (300 Å pore size, 5 μm particle size, 150 μm i.d. × 150 mm) at a flow rate of 1 μL/min. Peptides were separated with a gradient of 10–60% buffer B over 45 min, with buffer A (1% acetonitrile and 0.1% formic acid) and buffer B (80% acetonitrile with 0.1% formic acid). Gas and voltage settings were adjusted as required. Data Dependent Acquisition (DDA) was performed using an MS-TOF scan from an *m/z* of 350-1800 for 0.5 s followed by DDA MS/MS with automated selection of the top 20 peptides from an *m/z* of 350-1800 for 0.05 s per spectrum. Data Independent Acquisition (DIA) SWATH–MS was performed using identical LC conditions to DDA experiments, and with an MS–TOF scan from an *m/z* of 350-1800 for 0.05 s, followed by high-sensitivity information-independent acquisition with 26 *m/z* isolation windows, with 1 *m/z* window overlap each for 0.1 s across an *m/z* range of 400–1250. Collision energy was automatically assigned for each peptide by the Analyst software (SCIEX).

### Data analysis

Proteins were identified from DDA data using ProteinPilot 4.1 (SCIEX). The ID search settings were: sample type, identification; cysteine alkylation, acrylamide; instrument, TripleTof5600; species, human; ID focus, biological modifications; digestion, trypsin; Search effort, thorough ID. False discovery rate was conducted with limits of 99% confidence and 1% local false discovery rate, where peptides with a confidence greater than 99% were included in further analysis. An ion library from proteins identified with ProteinPilot was used to measure peptide abundance in each sample using PeakView 2.1 (SCIEX), with settings: shared peptides, allowed; peptide confidence threshold, 99%; false discovery rate, 1%; XIC extraction window, 6 min; XIC width, 75 ppm. The mass spectrometry proteomics data have been deposited to the ProteomeXchange Consortium via the PRIDE ^22^ partner repository with the dataset identifier PXD011331. Statistical analyses were performed as previously described ^23^ using ReformatMS ^24^ and MSstats (2.4) ^25^, a package for the R-software environment. Heat maps were prepared with GraphPad Prism and Venn diagrams with Venny 2.1 (http://bioinfogp.cnb.csic.es/tools/venny/). Hierarchical clustering was performed on log2-transformed normalized abundances of proteins with significant differences in abundance between groups with Cluster 3.0 using the average linkage method and Pearson’s correlation method for similarity. Trees were generated using JavaTreeview 1.1.6.

### Functional Enrichment analysis

Differentially abundant proteins (p < 10^-5^) were subjected to gene ontology (GO) term enrichment analysis using DAVID Bioinformatics resources ^26,27^ Significance was determined with a Benjamini-Hochberg adjusted p-value of < 0.05. The list of differentially abundant proteins that were identified in the AUD pre-frontal lobe was analysed using the protein database STRINGv10 database (http://string-db.org) to generate protein-protein interaction scores using default parameters ^28,29^. The protein-protein interaction scores were downloaded and transferred to Cytoscape for visualisation ^30^.

## Results

Pathophysiological sex-related differences in AUD patients have been previously reported ^31,32^, and drinking behaviour varies between ethnic groups ^33^. We therefore chose to exclusively study the effects of AUD in males of Northern European descent to limit confounding effects. We chose to study whole-tissue extracts as a means to glean generalised insights into the proteomes of diverse cell types and tissues.

### Protein Identification

We identified proteins from DDA LC-MS/MS of high pH fractionated pooled samples and unfractionated individual samples of trypsin-digested whole tissue protein extractions from pre-frontal lobe and motor cortex from human brain of AUD patients and controls. We then used SWATH-MS to measure the abundance of proteins in each brain region in each individual. A total of 614 unique quantifiable proteins were identified and measured with a critical local false discovery rate of < 0.01 (Supplementary Table 1).

### AUD has a profound effect on the overall soluble brain proteome

It has been proposed that molecular and tissue changes caused by AUD occur primarily in the pre-frontal lobe, with the motor cortex remaining comparatively unaffected ^34,35^. We therefore compared the proteome of the pre-frontal lobe in AUD patients and controls, and also compared the proteome of the motor cortex in AUD patients and controls. In addition, we compared the proteome of the pre-frontal lobe and the motor cortex in both controls and in AUD patients. All comparisons showed large and significant differences in abundance for a subset of the quantifiable proteins (Fig. 1, Supplementary Tables 2 - 5).

**Figure 1.**
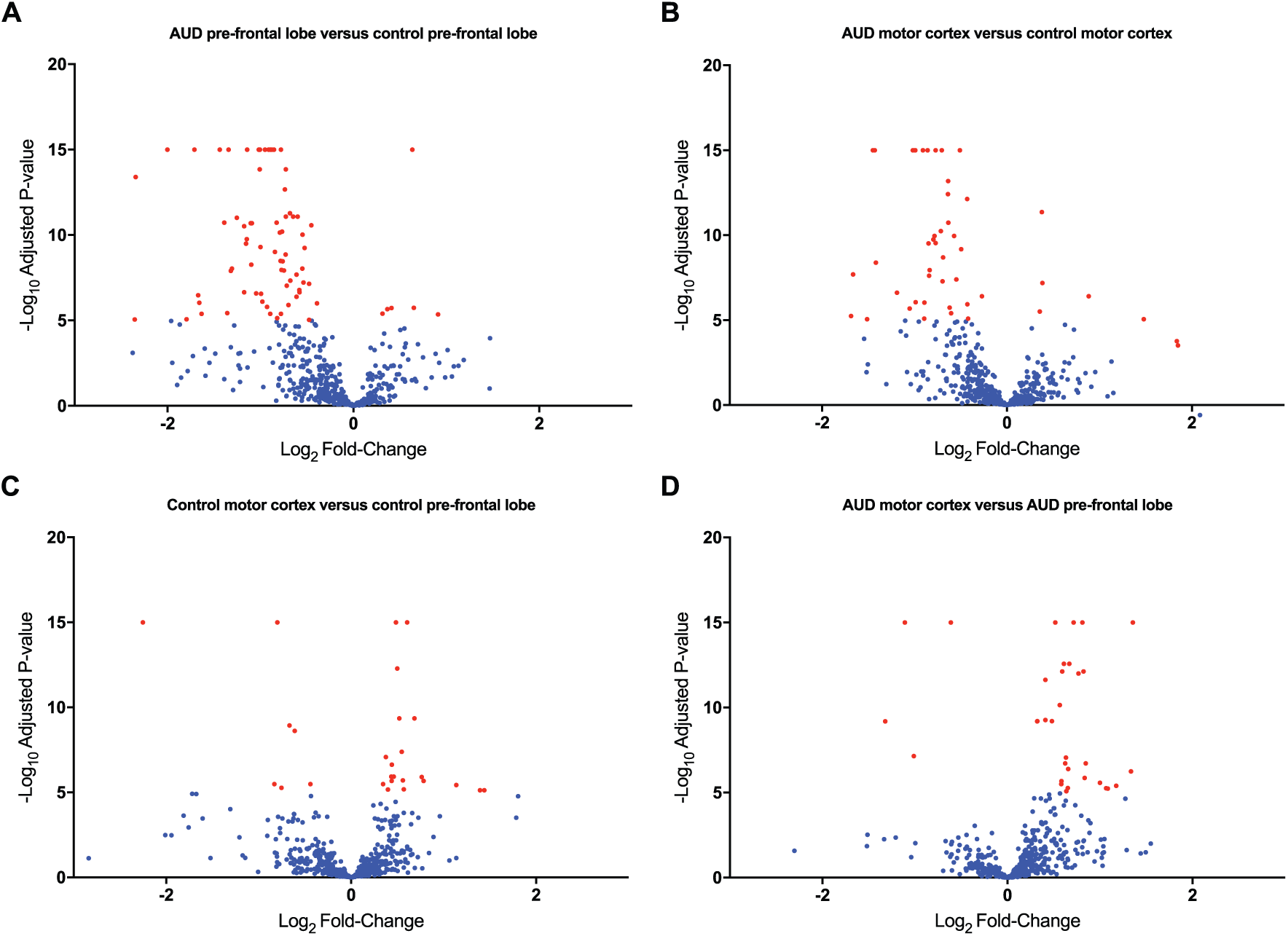
The pre-frontal lobe and motor cortex proteomes differ in AUD patients and controls. Volcano plots illustrating the comparisons between **(A)** AUD and control pre-frontal lobe, **(B)** AUD and control motor cortex, **(C)** motor cortex and pre-frontal lobe of controls, and **(D)** motor cortex and pre-frontal lobe of AUD patients. Red, significantly different in abundance (p < 10^-5^). Blue, not significantly different in abundance.

The proteome of the pre-frontal lobe showed major differences in AUD, with 70 proteins having significantly lower, and six proteins significantly higher abundance than in the controls (Fig. 1A, Supplementary Table 2). The motor cortex also differed substantially, with 50 and seven proteins detected with significantly higher and lower abundances than in controls, respectively (Fig. 1B, Supplementary Table 3). Statistically significant differences in both of these comparisons appear to be skewed towards proteins with lower abundance in AUD patients, suggesting AUD causes a reduction in the abundance of selected proteins in both the pre-frontal lobe and motor cortex. However, we note that our proteomics workflow measures relative, not absolute, protein abundance. We also observed differences in the proteomes of the pre-frontal lobe and motor cortex in both controls and AUD patients (Fig. 2C, D, Supplementary Table 3, 4). We expected to detect differences in these proteomes, as these brain areas are responsible for different cognitive and physiological processes. Surprisingly, it appeared that AUD had a larger effect on the proteome than brain region (Fig. 2A). To investigate this in more detail, we performed hierarchical clustering on the 126 proteins which showed statistically significant differences in abundance between AUD and controls or between brain regions. This analysis showed that the data clustered first on the basis of AUD patients versus controls, and next between pre-frontal lobe and motor cortex (Fig. 2). Together, this showed that AUD had a profound effect on the overall soluble brain proteome exceeding the inherent differences between brain regions.

**Figure 2.**
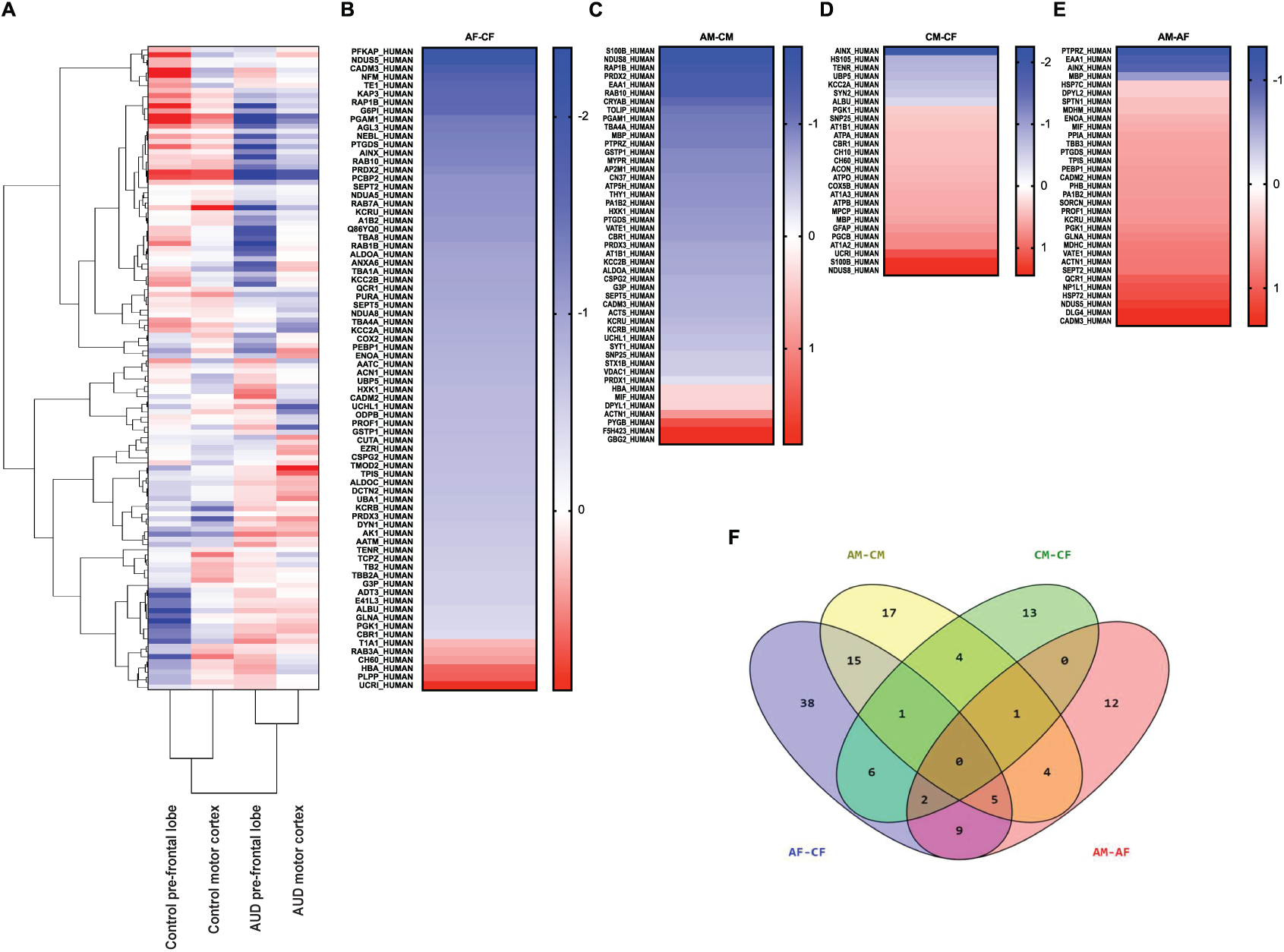
Proteomic differences associated with AUD are larger than between brain regions. **(A)** Hierarchical clustering and heat map of Log2-transformed normalised abundances of proteins with significantly different abundance. Blue, proteins decreased in abundance. Red, proteins increased in abundance. White, not significantly different. Vertical scales show the similarity between differentially abundant proteins, while horizontal scales show the similarity between tissue and disease state. Full data are shown in Supplementary Table 2. Heat maps of proteins with significant differences in abundance between **(B)** AUD and control pre-frontal lobe, **(C)** AUD and control motor cortex, **(D**) motor cortex and pre-frontal lobe of controls, and **(E)** pre-frontal lobe and motor cortex of AUD patients. **(F)** Venn diagram of proteins with significantly different abundance in each comparison. Full data are shown in Supplementary Table 6.

### Pathway analysis identifies differences in central metabolism in AUD patients

To discover higher-order pathways and processes altered in the brains of AUD patients, we performed GO term enrichment analysis on proteins that were significantly different in abundance. This showed that proteins in the pre-frontal lobe with lower abundance in AUD patients were strongly associated with glycolysis and gluconeogenesis (P < 0.05). No significant GO term enrichments were found in any other comparison. To identify additional molecular processes altered in the brains of AUD patients, we performed pathway analysis using STRING, and obtained an interaction map of proteins with significant differences in abundance in the AUD pre-frontal lobe (Fig. 3). This interaction map showed strong connectivity for proteins involved in glycolysis (PFKAP and G3P), with additional links to proteins of the Ras superfamily (RAP1B and RAB10), and proteins involved in oxidative phosphorylation (NDUS8 and AATC), the cytoskeleton (TBA8), and neurotoxicity (PRDX2 and AATM) (Fig. 4).

As our cohort of AUD patients was older than the controls, we performed correlation analysis of subject age and normalised protein abundance for each selected protein that was significantly different in abundance between AUD patients and controls. None of the proteins in the pre-frontal lobe with significantly different abundance in AUD compared to controls were found to be correlated with age (Supplementary Table 7), confirming that the differences in protein abundance we observed in the pre-frontal lobe were dependent on AUD status. Some age-dependent changes in protein abundance were observed in the motor cortex (Supplementary Table 7).

**Figure 3.**
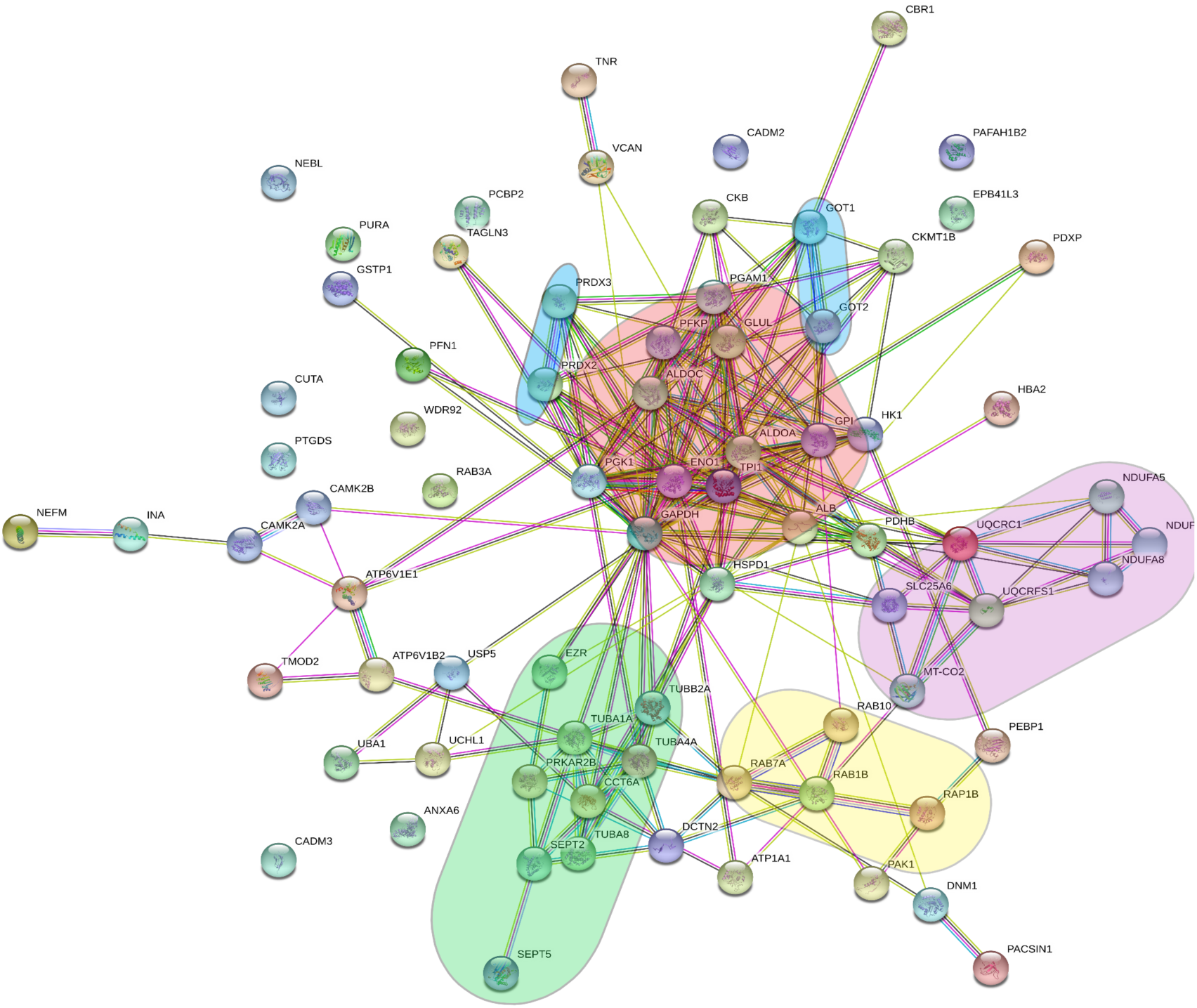
STRING protein interaction map of differentially abundant proteins in the AUD pre-frontal lobe. Protein hubs are highlighted: red, glycolysis; purple, oxidative phosphorylation; yellow, Ras superfamily; green, cytoskeleton; blue, excitotoxicity.

**Figure 4.**
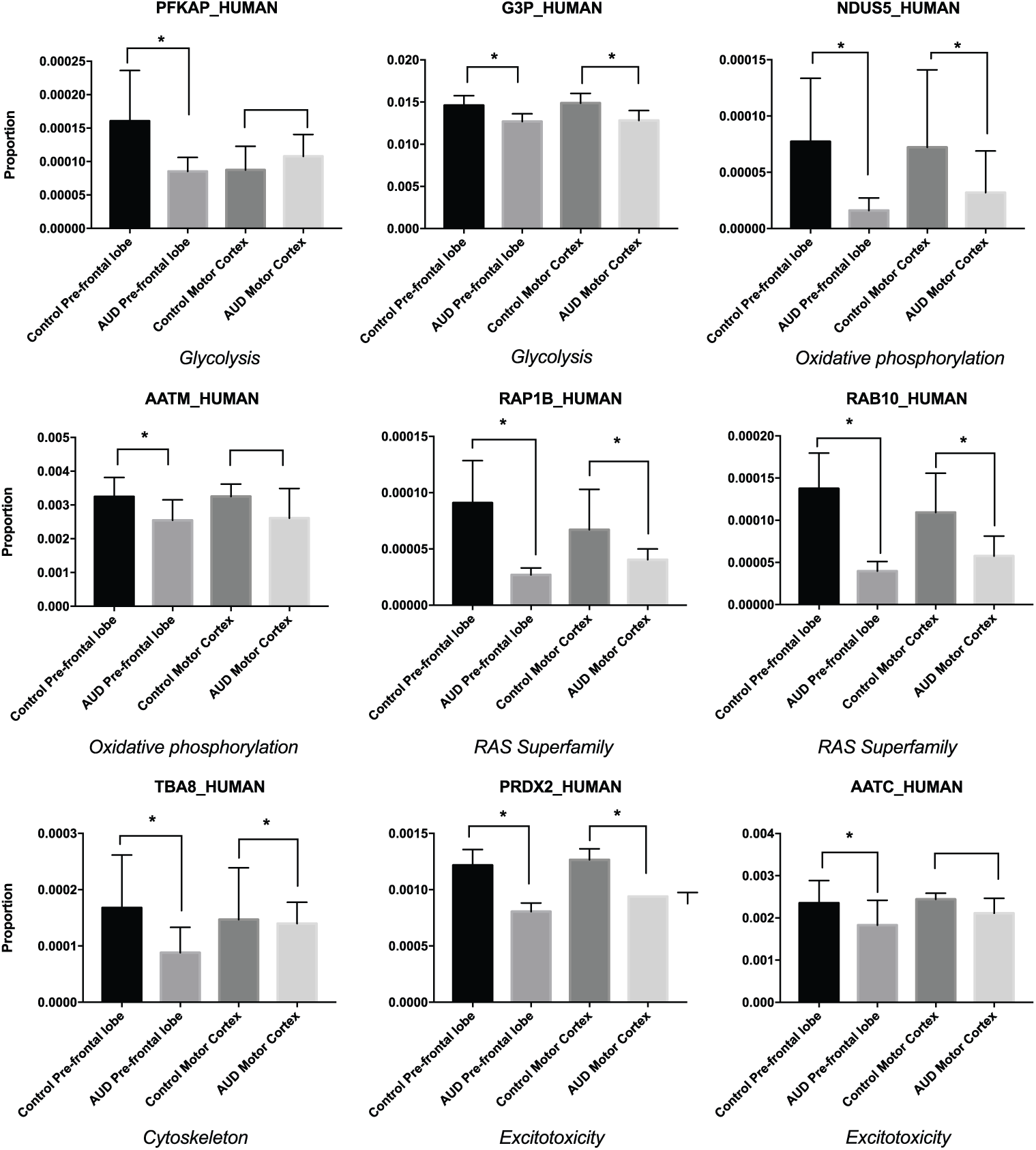
Abundance of selected proteins of interest in the AUD pre-frontal lobe. Proteins were selected according to their function and distance from the central glycolytic hub. *, significant difference in abundance, p < 10^-5^

## Discussion

We used SWATH-MS to compare the proteomes of pre-frontal lobe and motor cortex of AUD patients and controls. While we detected differences in the proteomes of pre-frontal lobe and motor cortex in both controls and AUD patients, the AUD-dependent changes in region-specific proteomes were substantially larger (Fig. 1 and 2). This emphasizes the profound effects on brain physiology associated with AUD.

### Alcohol use disorder perturbs central brain metabolism

GO term enrichment analysis enabled us to identify nine enzymes involved in glycolysis that were less abundant in AUD patients in the pre-frontal lobe. Many of these proteins were also less abundant in AUD patients in the motor cortex. Phosphofructokinase (PFKAP), the first committed step in glycolysis ^36^, was the protein with the largest decrease in abundance in the pre-frontal lobe in AUD compared to controls (Fig. 2 and 4, Supplementary Table 2). Several other glycolytic enzymes were also less abundant in AUD patients, including the pyruvate dehydrogenase complex, which catalyses the production of acetyl-CoA from pyruvate required for the TCA cycle ^37^ Together, our data suggest that there is a substantial reduction in glycolytic flux in AUD. This is consistent with previous findings in human imaging studies that indicated that low doses of alcohol significantly decrease baseline brain glucose metabolism ^38,39^, and that alcohol misuse causes a reduction in glucose metabolism ^40^.

Alcohol is primarily metabolized in the cytosol of hepatocytes by alcohol dehydrogenase to acetaldehyde ^41^, which is further oxidized to acetate ^42^. This excess acetate then enters the blood circulation. Previous studies have reported an increased uptake of acetate into the alcoholic brain, where it is metabolised in astrocytes to acetyl-CoA to enter the TCA cycle ^43^. While our proteomic data identified all enzymes of the TCA cycle, none of these showed significant changes in abundance in AUD (Supplementary Table 2). Consumption of acetyl-CoA in the TCA cycle results in a net production of NADH ^44^ Increased acetate metabolism in the AUD brain would therefore be expected to increase the local NADH:NAD+ ratio. This affects many metabolic processes, and in particular glycolysis, which requires NAD+. Together, our data are consistent with AUD being associated with (i) a switch in energy utilisation in the brain from glucose towards acetate, (ii) enhanced metabolism of acetate through the TCA cycle, (iii) an increase in cellular NADH, with (iv) a concomitant severe down-regulation of glycolysis. These changes in cellular metabolism may contribute to alcohol dependence through a shift to a metabolic preference for ethanol-derived acetate.

### Microtubules, vesicular trafficking, and the cytoskeleton

GTP-binding proteins from the RAS superfamily are critical components of diverse intracellular signalling networks ^45^. Rap-1b and Rab-10 were significantly less abundant in both the AUD pre-frontal lobe and motor cortex, while Rab-7a and Rab-1b were significantly less abundant in the AUD pre-frontal lobe (Fig. 4, Supplementary Table 2). Rap-1b is a GABAβ receptor binding partner, responsible for initiating the signalling cascade that mediates GABAβ receptor cell surface expression via receptor recycling ^46^. Normal functioning of Rap-1b results in slow and long-lasting inhibitory action via the GABA neurotransmitter ^46^. The observation that Rap-1b was less abundant in AUD in both brain areas suggests that the AUD brain has a reduced capacity to modulate inhibitory outputs, in particular in the mesocorticolimbic dopamine system responsible for addiction ^47^ GABAβ is a potential treatment target for AUD, as demonstrated by the use of the receptor agonist baclofen in reducing craving and withdrawal symptoms ^48^. Rab-10 is proposed to be involved in neurite outgrowth during the processes of axonal growth, pathfinding, regeneration, and dendritic growth ^49^. These processes require extensive membrane trafficking events for normal functioning^50-52^. Reduction in such intrinsic axon growth ability may suggest that the AUD brain is deficient in integration of synaptic inputs. The retromer is a protein complex associated with endosome to trans-Golgi network trafficking ^53,54^ Rab-7a is a retromer effector responsible for cargo-recognition for the retrograde pathway for transmembrane receptor recycling^55^. Down-regulation of Rab-7a could therefore disrupt transmembrane receptor recycling to the pre- and post-synaptic membranes, adversely affecting the delicate neurochemistry of the pre-frontal lobe.

Microtubules are a fundamental cytoskeletal component of neurons that are important in a multitude of cellular and developmental processes. The core microtubule structure consists of heterodimers of α and β tubulin isoforms. Our data showed that tubulin alpha chains (TUBA1A, TUBA4A, and TUBA8) were less abundant in AUD in both the pre-frontal lobe and motor cortex, and that beta chain TUBB2A was less abundant in AUD in the pre-frontal lobe (Fig. 4, Supplementary Table 2). Loss of cytoskeletal protein is associated with selective neuronal tissue loss and tissue pathology observed in alcoholics ^34^

### Excitotoxicity and oxidative damage in the AUD brain

Excitotoxicity is a common pathological hallmark of neurodegenerative diseases. This process results in a calcium overload in the mitochondria, triggering compensatory respiration and the generation of reactive oxygen species (ROS) ^56,57^ Long-term ROS irreversibly damage the electron transport chain and initiate apoptotic and necrotic events within the cell ^58^. Two members of the antioxidant peroxiredoxin family effective against hydrogen peroxide and peroxynitrite, peroxiredoxin 2 and 3, were significantly decreased in abundance in both the pre-frontal lobe and the motor cortex in patients with AUD. Enzyme kinetic experiments have demonstrated that hyperoxidation of peroxiredoxin 1 causes rapid inactivation of the enzyme, a phenomenon which is conserved in peroxiredoxin 2 and 3 ^59,60^. Peroxiredoxin 2 is more susceptible to hyperoxidation due to slower disulphide formation ^61^. This is consistent with our data, as peroxiredoxin 3 had a smaller AUD-dependent fold-change than peroxiredoxin 2 in both brain locations. Our data are therefore consistent with increased oxidative stress in AUD, causing chronic overoxidation and degradation of peroxiredoxins.

A key example of neuro-excitotoxicity is glutamate toxicity ^62^. Chronic alcoholism has been associated with hyperglutamatergic activity in the mesocorticolimbic brain areas ^63^. Alcohol non-competitively inhibits NMDA receptors, the binding targets of glutamate ^64^ The enzymes cytoplasmic aspartate aminotransferase (AATC; also known as GOT1) and glutamine synthetase (GLUL) are important regulators of glutamate levels and have been implicated in brain neuroprotection ^65^. Our data showed a decrease in AATC and GLUL abundance in the AUD pre-frontal lobe (Fig. 4), consistent with increased levels of glutamate and enhanced excitotoxity in the AUD brain.

### Epigenetic and post-translational modifications

The differences we observed in the proteomes in the brains of subjects with AUD may be associated with epigenetic regulation and/or protein post-translational modifications (PTM) ^66^. Epigenetic changes have previously been implicated in alcohol abuse, where changes in NADH:NAD^+^ ratios affect the activity of proteins responsible for gene expression and silencing ^67,68^. Increased DNA methylation has been associated with a decrease in synaptic proteins in the pre-frontal lobe of rats with alcohol-induced behaviour^69^. In addition, acetyl-CoA is used in histone acetylation by histone acetyltransferases to regulate gene expression ^70^, and also modifies diverse other proteins with acetylation to regulate their functions. The PTM of acetylation is dynamic and reversible, but changes in the NADH:NAD^+^ ratio associated with AUD would be expected to perturb the balance of this modification. Alcohol-induced acetylation has been shown to affect several proteins ^71^. GAPDH has emerged as an important regulator of metabolism, with acetylation at multiple lysine residues potentially affecting its activity ^72^. It has also been reported that lysine acetylation of *α* tubulins is increased in alcoholic cases compared with controls ^34^ It is therefore possible that an increased NADH:NAD^+^ ratio and acetyl-CoA concentration in the AUD brain cause inappropriate increases in DNA methylation and protein acetylation that systemically perturb translation and directly disrupt the functions of diverse proteins.

## Conclusion

Using SWATH-MS proteomic analyses we identified large and significant AUD-dependent differences in the abundance of proteins in pre-frontal lobe and motor cortex. We observed profound changes in metabolic enzymes consistent with a switch from glucose to acetate utilization in the AUD brain. The proteins with differences in abundance in AUD may be useful as biomarkers of disease. Further studies are necessary to clarify the mechanisms underlying these differences, whether they are causes or effects of AUD, and the potential role of DNA methylation and/or lysine acetylation in regulating these changes. We also propose that metabolic adaptations allowing efficient acetate utilization contribute to ethanol dependence in AUD.

## Acknowledgements

We gratefully acknowledge the assistance of Dr Amanda Nouwens and Mr Peter Josh at The University of Queensland School of Chemistry and Molecular Biosciences Mass Spectrometry Facility. Benjamin L. Schulz is funded by an NHMRC Career Development Fellowship APP1087975. Edward D. Kerr is funded by an Advance Queensland PhD Scholarship.

## References

1 World Health Organisation. Global strategy to reduce harmful use of alcohol. (2010).

2 Oscar-Berman, M. & Bowirrat, A. Genetic influences in emotional dysfunction and alcoholism-related brain damage. Neuropsychiatr Dis Treat 1, 211–229, (2005).

3 Oscar-Berman, M., Kirkley, S. M., Gansler, D. A. & Couture, A. Comparisons of Korsakoff and non-Korsakoff alcoholics on neuropsychological tests of prefrontal brain functioning. Alcohol Clin Exp Res 28, 667–675, (2004).

4 Lewohl, J. M. et al. Gene expression in human alcoholism: microarray analysis of frontal cortex. Alcohol Clin Exp Res 24, 1873–1882, (2000).

5 Erickson, E. K., Farris, S. P., Blednov, Y. A., Mayfield, R. D. & Harris, R. A. Astrocyte-specific transcriptome responses to chronic ethanol consumption. Pharmacogenomics J 18, 578–589, (2018).

6 Farris, S. P., Arasappan, D., Hunicke-Smith, S., Harris, R. A. & Mayfield, R. D. Transcriptome organization for chronic alcohol abuse in human brain. Mol Psychiatry 20, 1438–1447, (2015).

7 de la Monte, S. M. & Kril, J. J. Human alcohol-related neuropathology. Acta Neuropathol 127, 71–90, (2014).

8 Vargas, W. M., Bengston, L., Gilpin, N. W., Whitcomb, B. W. & Richardson, H. N. Alcohol binge drinking during adolescence or dependence during adulthood reduces prefrontal myelin in male rats. J Neurosci 34, 14777–14782, (2014).

9 Liu, Y., Beyer, A. & Aebersold, R. On the Dependency of Cellular Protein Levels on mRNA Abundance. Cell 165, 535–550, (2016).

10 Doyle, J. P. et al. Application of a translational profiling approach for the comparative analysis of CNS cell types. Cell 135, 749–762, (2008).

11 Sugino, K. et al. Molecular taxonomy of major neuronal classes in the adult mouse forebrain. Nat Neurosci 9, 99–107, (2006).

12 Warden, A. S. & Mayfield, R. D. Gene expression profiling in the human alcoholic brain. Neuropharmacology 122, 161–174, (2017).

13 Gorini, G., Adron Harris, R. & Dayne Mayfield, R. Proteomic Approaches and Identification of Novel Therapeutic Targets for Alcoholism. Neuropsychopharmacology 39, 104–130, (2014).

14 Alexander-Kaufman, K., James, G., Sheedy, D., Harper, C. & Matsumoto, I. Differential protein expression in the prefrontal white matter of human alcoholics: a proteomics study. Mol Psychiatry 11, 56–65, (2006).

15 Kashem, M. A., Harper, C. & Matsumoto, I. Differential protein expression in the corpus callosum (genu) of human alcoholics. Neurochem Int 53, 1–11, (2008).

16 Matsumoto, I. Proteomics approach in the study of the pathophysiology of alcohol-related brain damage. Alcohol Alcohol 44, 171–176, (2009).

17 Chang, R. Y., Etheridge, N., Dodd, P. R. & Nouwens, A. S. Targeted quantitative analysis of synaptic proteins in Alzheimer’s disease brain. Neurochem Int 75, 66–75, (2014).

18 Zhuang, A. et al. Increased liver AGEs induce hepatic injury mediated through an OST48 pathway. Sci Rep 7, 12292, (2017).

19 Nguyen, L. T., Zacchi, L. F., Schulz, B. L., Moore, S. S. & Fortes, M. R. S. Adipose tissue proteomic analyses to study puberty in Brahman heifers. JAnim Sci 96, 2392–2398, (2018).

20 Ni, M. W. et al. Modified filter-aided sample preparation (FASP) method increases peptide and protein identifications for shotgun proteomics. Rapid Commun Mass Spectrom 31, 171–178, (2017).

21 Xu, Y., Bailey, U. M. & Schulz, B. L. Automated measurement of site-specific N-glycosylation occupancy with SWATH-MS. Proteomics 15, 2177–2186, (2015).

22 Vizcaíno, J. A. et al. 2016 update of the PRIDE database and its related tools. Nucleic Acids Research 44, D447–D456, (2016).

23 Zacchi, L. F. & Schulz, B. L. SWATH-MS Glycoproteomics Reveals Consequences of Defects in the Glycosylation Machinery. Mol Cell Proteomics 15, 2435–2447, (2016).

24 Kerr, E. D., Phung, T. K., Fox, G. P., Platz, G. J. & Schulz, B. L. The intrinsic and regulated proteomes of barley seeds in response to fungal infection. bioRxiv, (2018).

25 Choi, M. et al. MSstats: an R package for statistical analysis of quantitative mass spectrometry-based proteomic experiments. Bioinformatics 30, 2524–2526, (2014).

26 Huang da, W., Sherman, B. T. & Lempicki, R. A. Bioinformatics enrichment tools: paths toward the comprehensive functional analysis of large gene lists. Nucleic Acids Res 37, 1–13, (2009).

27 Huang da, W., Sherman, B. T. & Lempicki, R. A. Systematic and integrative analysis of large gene lists using DAVID bioinformatics resources. Nature protocols 4, 44–57, (2009).

28 Szklarczyk, D. et al. STRING v10: protein-protein interaction networks, integrated over the tree of life. Nucleic Acids Res 43, D447–452, (2015).

29 Szklarczyk, D. et al. The STRING database in 2017: quality-controlled protein-protein association networks, made broadly accessible. Nucleic Acids Res 45, D362–d368, (2017).

30 Shannon, P. et al. Cytoscape: a software environment for integrated models of biomolecular interaction networks. Genome Res 13, 2498–2504, (2003).

31 Barton, E. A., Baker, C. & Leasure, J. L. Investigation of Sex Differences in the Microglial Response to Binge Ethanol and Exercise. Brain Sci 7, (2017).

32 West, R. K., Maynard, M. E. & Leasure, J. L. Binge ethanol effects on prefrontal cortex neurons, spatial working memory and task-induced neuronal activation in male and female rats. Physiol Behav 188, 79–85, (2018).

33 Gelernter, J. et al. Genomewide Association Study of Alcohol Dependence and Related Traits in a Thai Population. Alcohol Clin Exp Res 42, 861–868, (2018).

34 Erdozain, A. M. et al. Alcohol-related brain damage in humans. PloS one 9, e93586, (2014).

35 Kril, J. J., Halliday, G. M., Svoboda, M. D. & Cartwright, H. The cerebral cortex is damaged in chronic alcoholics. Neuroscience 79, 983–998, (1997).

36 Berg, J. M., Tymockzo, J. L. & Stryer, L. in Biochemistry (W.H. Freeman, 2002).

37 Patel, M. S., Nemeria, N. S., Furey, W. & Jordan, F. The pyruvate dehydrogenase complexes: structure-based function and regulation. J Biol Chem 289, 16615–16623, (2014).

38 Rae, C. D. et al. Ethanol, not metabolized in brain, significantly reduces brain metabolism, probably via specific GABA(A) receptors. J Neurochem 129, 304–314, (2014).

39 Volkow, N. D. et al. Alcohol Decreases Baseline Brain Glucose Metabolism More in Heavy Drinkers Than Controls But Has No Effect on Stimulation-Induced Metabolic Increases. J Neurosci 35, 3248–3255, (2015).

40 Volkow, N. D. et al. Low doses of alcohol substantially decrease glucose metabolism in the human brain. Neuroimage 29, 295–301, (2006).

41 Agarwal, D. P. & Goedde, H. W. Human aldehyde dehydrogenases: their role in alcoholism. Alcohol 6, 517–523, (1989).

42 Eriksson, C. J. The role of acetaldehyde in the actions of alcohol (update 2000). Alcohol Clin Exp Res 25, 15s–32s, (2001).

43 Jiang, L. et al. Increased brain uptake and oxidation of acetate in heavy drinkers. J Clin Invest 123, 1605–1614, (2013).

44 Cederbaum, A. I. Alcohol metabolism. Clin Liver Dis 16, 667–685, (2012).

45 Goitre, L., Trapani, E., Trabalzini, L. & Retta, S. F. The Ras superfamily of small GTPases: the unlocked secrets. Methods Mol Biol 1120, 1–18, (2014).

46 Zhang, Z. et al. GABAB receptor promotes its own surface expression by recruiting a Rap1-dependent signaling cascade. J Cell Sci 128, 2302–2313, (2015).

47 Jayaram, P. & Steketee, J. D. Effects of repeated cocaine on medial prefrontal cortical GABAB receptor modulation of neurotransmission in the mesocorticolimbic dopamine system. J Neurochem 90, 839–847, (2004).

48 Agabio, R. & Colombo, G. GABAB receptor ligands for the treatment of alcohol use disorder: preclinical and clinical evidence. Front Neurosci 8, (2014).

49 Chua, C. E. L. & Tang, B. L. Rab 10-a traffic controller in multiple cellular pathways and locations. J Cell Physiol, (2018).

50 Bloom, O. E. & Morgan, J. R. Membrane trafficking events underlying axon repair, growth, and regeneration. Mol Cell Neurosci 48, 339–348, (2011).

51 Meldolesi, J. Neurite outgrowth: this process, first discovered by Santiago Ramon y Cajal, is sustained by the exocytosis of two distinct types of vesicles. Brain Res Rev 66, 246–255, (2011).

52 Pfenninger, K. H. Plasma membrane expansion: a neuron’s Herculean task. Nat Rev Neurosci 10, 251–261, (2009).

53 Burd, C. & Cullen, P. J. Retromer: A Master Conductor of Endosome Sorting. Cold Spring Harb Perspect Biol 6, a016774, (2014).

54 Seaman, M. N. J. Recycle your receptors with retromer. Trends Cell Biol 15, 68–75, (2005).

55 Harrison, M. S. et al. A mechanism for retromer endosomal coat complex assembly with cargo. Proc Natl Acad Sci U S A 111, 267–272, (2014).

56 Hughes, J. R. Alcohol withdrawal seizures. Epilepsy Behav 15, 92–97, (2009).

57 Stavrovskaya, I. G. & Kristal, B. S. The powerhouse takes control of the cell: Is the mitochondrial permeability transition a viable therapeutic target against neuronal dysfunction and death? Free Radic Biol Med 38, 687–697, (2005).

58 Hattori, F., Murayama, N., Noshita, T. & Oikawa, S. Mitochondrial peroxiredoxin-3 protects hippocampal neurons from excitotoxic injury in vivo. J Neurochem 86, 860–868, (2003).

59 Perkins, A., Nelson, K. J., Parsonage, D., Poole, L. B. & Karplus, P. A. Peroxiredoxins: guardians against oxidative stress and modulators of peroxide signaling. Trends Biochem Sci 40, 435–445, (2015).

60 Yang, K. S. et al. Inactivation of human peroxiredoxin I during catalysis as the result of the oxidation of the catalytic site cysteine to cysteine-sulfinic acid. J Biol Chem 277, 38029–38036, (2002).

61 Peskin, A. V. et al. Hyperoxidation of peroxiredoxins 2 and 3: rate constants for the reactions of the sulfenic acid of the peroxidatic cysteine. J Biol Chem 288, 14170–14177, (2013).

62 Lewerenz, J. & Maher, P. Chronic Glutamate Toxicity in Neurodegenerative Diseases-What is the Evidence? Front Neurosci 9, 469, (2015).

63 Zou, J. et al. Glutamine synthetase down-regulation reduces astrocyte protection against glutamate excitotoxicity to neurons. Neurochem Int 56, 577–584, (2010).

64 Rao, P. S., Bell, R. L., Engleman, E. A. & Sari, Y. Targeting glutamate uptake to treat alcohol use disorders. Front Neurosci 9, 144, (2015).

65 Ruban, A. et al. Combined Treatment of an Amyotrophic Lateral Sclerosis Rat Model with Recombinant GOT1 and Oxaloacetic Acid: A Novel Neuroprotective Treatment. Neurodegener Dis 15, 233–242, (2015).

66 Zakhari, S. Alcohol metabolism and epigenetics changes. Alcohol Res 35, 6–16, (2013).

67 Imai, S., Armstrong, C. M., Kaeberlein, M. & Guarente, L. Transcriptional silencing and longevity protein Sir2 is an NAD-dependent histone deacetylase. Nature 403, 795–800, (2000).

68 Kumar, V. et al. Transcription corepressor CtBP is an NAD(+)-regulated dehydrogenase. Mol Cell 10, 857–869, (2002).

69 Barbier, E. et al. DNA methylation in the medial prefrontal cortex regulates alcohol-induced behavior and plasticity. J Neurosci 35, 6153–6164, (2015).

70 Lee, K. K. & Workman, J. L. Histone acetyltransferase complexes: one size doesn’t fit all. Nat Rev Mol Cell Biol 8, 284–295, (2007).

71 Shepard, B. D. & Tuma, P. L. Alcohol-induced protein hyperacetylation: mechanisms and consequences. World J Gastroenterol 15, 1219–1230, (2009).

72 Bond, S. T. et al. Lysine post-translational modification of glyceraldehyde-3-phosphate dehydrogenase regulates hepatic and systemic metabolism. FASEB J 31, 2592–2602, (2017).

